# Characterisation of gastrointestinal helminths and their impact in commercial small-scale chicken flocks in the Mekong Delta of Vietnam

**DOI:** 10.1101/628024

**Authors:** Nguyen T.B. Van, Nguyen V Cuong, Nguyen T. Phuong Yen, Nguyen T.H. Nhi, Bach Tuan Kiet, Nguyen V. Hoang, Vo B. Hien, Guy Thwaites, Juan J. Carrique-Mas, Alexis Ribas

## Abstract

Commercial small-scale chicken farms using all-in-all-out production but operating with low standards of hygiene/biosecurity are increasingly common in Vietnam. These conditions facilitate transmission of gastrointestinal helminths. However there are no published data on these parasites in these systems. The aims were: (1) to determine the prevalence/burden of gastrointestinal helminths in small-scale commercial flocks in commercial small-scale flocks in the Mekong Delta region; and (2) to investigate the association between worm burdens and birds’ weight and disease status. Randomly selected chickens (n=120) (‘normal’ flocks) were investigated at the end of their production cycle (∼18 weeks), as well as 90 chickens with signs of respiratory and/or severe disease. The gastrointestinal tract of chickens was dissected and all visible helminths were identified. 54.2% and 54.4% healthy and diseased chickens contained helminths. Diseased, colonized chickens harboured a higher mass of helminth worms (3.8 ±SD 8.6g) than colonized, healthy chickens (1.9 ±6.3g). Eight species were identified, three nematodes (*Ascaridia galli, Cheilospirura hamulosa* and *Heterakis gallinarum*), four cestodes (*Hymenolepis, Raillietina cesticillus, Raillietina echinobothrida, Raillietina tetragona*,) and one trematode (Echinostomatidae). *Heterakis gallinarum* was the most prevalent helminth (43.3% and 42.2% in healthy and sick chickens, respectively), followed by *A. galli* (26.7% and 41.1%). Colonized chickens weighed 101.5g less than non-colonized birds. Colonisation was significantly higher during the rainy months (May-November) for both *H. gallinarum* and *A. galli*. Anthelminthic usage was not associated with reduced helminth burdens. We recommend upgrading cleaning and disinfection and limiting access to ranging areas to control helminth infections in small-scale commercial chicken flocks.

## Introduction

Gastrointestinal helminth parasites represent a major constraint to the productivity of small-scale and backyard poultry farming worldwide (Swayne 2013). These helminths interfere with the host’s metabolism, resulting in poor feed utilization, reducing growth rate and productivity (Gauly et al. 2007, Phiri et al. 2007). In addition, helminths increase disease susceptibility and impair the immunological response to vaccination (Kunchara Na Ayudthaya and Sangvaranond 1997, Pleidrup et al. 2014). Furthermore, gastrointestinal helminths can transmit pathogenic agent such as *Histomonas meleagridis*, which causes high morbidity and up to 20 % mortality in chicken flocks (McDougald 2005).

The Mekong Delta hosts about a fifth of Vietnam’s poultry population (VCNST 2018), and poultry production engages ∼50% rural households (Lan Phuong et al. 2015). Most of the chicken production is non-intensive, and occurs in mixed farms where chickens are raised in backyards, gardens, and orchards. However over recent years, many Mekong Delta farmers have been upgrading from backyard to confined ‘all-in-all-out’ systems using slow-growing local breeds. The number of farms raising more than 100 chickens in Vietnam has increased 41.5% 2011 to 2016 (Anon. 2018a). However there are no published studies on gastrointestinal helminths in chickens flocks in Vietnam. Studies from India and Thailand have evidenced a high (>70 %) prevalence of colonization with helminths in chickens (Butboonchoo and Wongsawad 2017, Katoch et al. 2012a, Kunchara Na Ayudthaya and Sangvaranond 1997, Yadav and Tandon 1991). However such studies were based on chickens collected in local markets, and factors that determine the assemblages of their helminth faunas were not investigated. We carried out a field survey aimed at caracterizing gastrointestinal helminths in ‘representative’ small-scale chicken flocks at the end of their production cycle, as well as in flocks presenting with severe symptoms of disease presenting to the veterinary authorities in the Mekong Delta province of Dong Thap. Specific objectives were: (1) to determine the prevalence and burden of gastrointestinal helminths in small-scale, confined flocks; and (2) to investigate the potential association between the burdens of helminth infection on the birds’ weight and disease status, as well as other farm, husbandry, and climatic variables.

## Methods

### Farms and study location

This study was carried out in Dong Thap, a province located in the Mekong Delta region (southwest of Vietnam). The province has a population density of ∼510 per km^2^ (Anon. 2014), and is dominated by flood plains. Their main agricultural outputs are rice, fruits, fisheries and livestock (ducks, chickens and pigs). The climate in the Mekong Delta is tropical, with an average temperature of 27.8°C, with little seasonal variation. Total annual rainfall typically ranges from 1,400 to 2,400mm, with a rainy season from May to November, accounting for ∼90% total rainfall. Farms raising between 100 and 2,000 birds raising chickens as single age flocks, randomly selected from the official provincial census, were enrolled into a study from two districts (Cao Lanh and Thap Muoi) within the study province between May 2017 and March 2018 (‘normal’ flocks) (Carrique-Mas and Rushton 2017). A television spot was run on local TV to ask owners of chicken flocks with signs of respiratory and/or severe disease to contact the provincial veterinary authorities (Sub-Department of Animal Health and Production of Dong Thap, SDAHP-DT) (‘diseased’ flocks). The study was conducted between September 2017 and March 2018.

### Sample and data collection

‘Normal’ flocks were visited at the end of their production cycle (typically 3-5 months) where one representative chicken was collected to be examined for the presence of helminths. In addition, data on the famer’s demographic characteristics, the flock, and husbandry practices were collected using structured questionaires. Data collected from flocks included: (1) Demographics of farmer (age, gender); (2) Highest level of education attained; (3) Experience in chicken farming (years); (4) Type of chicken house floor; (5) Density of chickens in house/pen (chickens/m^2^); (6) Presence of other poultry in the farm (other chicken flocks, ducks and muscovy ducks); (7) Season at the time of necropsy (‘rainy’ season, May to November, ‘dry’ season, December to April); (8) Number of chicks stocked; (9) Age of the flock (in weeks). The following data was only available from normal flocks: (10) Anthelminthics used over the production cycle; (11) Percent of weeks when farmers reported disease over the life of the flock (any disease, malaise, diarrhoea or respiratory signs); (12) Average weekly mortality over the production cycle. Data were entered into an Access (Microsoft Office) database. From each farm one (normal flock) or two (diseased flocks) representative chickens were selected to carry out a post-mortem investigation. The study was approved by the People’s Committee of the Province of Dong Thap. All farm visits were carried out by veterinarians affiliated to the SDAH-DT.

### Examination of chickens for the presence of helminths

Chickens were euthanized following humane procedures (Anon. 2013) by a trained veterinarian. The birds were weighed and their gastrointestinal tract (GIT) was examined for the presence of helminths. The GIT of each bird was systematically separated into three parts: (1) gizzard and proventriculus combined, (2) small intestine (duodenum, jejunum and ileum) and (3) caeca. The gizzard and proventriculus were dissected and examined using a binocular microscope. The small intestine was placed on a large Petri dish containing saline solution. A pair of scissors was used to dissect it and a scalpel to remove all worms seen attached to the mucosal surface; then the small intestine and its contents were further tranfered to a large (500 ml) plastic container, filled with tap water and shaken vigorously. These contents were then filtered several times using a sieve (mesh size 75 microns), and emptied onto a Petri dish. The two ceca were dissected in a Petri dish containing saline solution to facilitate detachment of worms from faeces and mucosa. All worms seen were transferred onto 70 % ethyl alcohol-containing tubes pre-labelled with the organ of collection and helminth type (nematode, cestode, trematode). All worms were counted and identified using a binocular microscope.

### Helminth species identification and DNA barcoding

Helminths were identified based on their morphological characteristics (Gomes et al. 2004, Soulsby 1982), and a subset of samples were used for molecular confirmation and identification using DNA barcoding (Gasser and Hoste 1995, Ribas et al. 2013). DNA extraction was performed using DNeasy Blood and Tissue kit (Qiagen, USA). The first and second internal transcribed spacers (ITS1 and ITS2) as well as the 5.8S rDNA gene (Gasser and Hoste 1995) were amplified. The PCR reaction used a conserved oligonucleotide primers NC5> 5’-GTAGGTGAACCTGCGGAAGGATCATT-3” (forward) and <NC2 5’-TTAGTTTCTTTTCCTCCGCT-3” (reverse). The thermal cycling profile included an initial denaturation in 94 °C for 3 min, followed by 40 cycles at 94°C for 45 sec, 53°C for 30 sec and 72°C for 1 min, and a final extension cycle at 72°C for 10 min. PCR products were separated on 1.4% agarose gel run in 1% TBE buffer under constant 120V for 1 h and the gel stained by Nancy-520 and visualized under UV light. The PCR products were amplified and sequenced using the Big Dye Cycle Sequencing kit (Applied Biosystems, USA) on an ABI 3770 automatic sequence. After generating a sequence type, sample was inferred to species according to the data available on the Basic Local Alignment Search Tool (BLAST) at the the National Center for Biotechnology Information (NCBI).

### Statistical analyses

A sample size of 120 normal chickens and 90 diseased chickens was chosen based on an expected prevalence of helminth colonization of 65% (in normal) and 80% (in diseased) flocks, respectively, estimated with a precision of 0.085 and a 95 % confidence level. The *χ*^2^ test was performed to compare the prevalence of helminths (total and by species) between ‘normal’ and ‘diseased’ flocks. The association between rainfall and prevalence of different helminth species was investigated. The agreement between presence/absence of different helminth species in the same bird, as well as over several cycles in the same farm was investigated by calculating the Kappa statistic. We transformed worm counts of each species to ‘mass of helminth worms’ based on the formula:

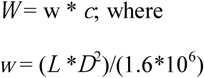

where *W* is the mass of helminth worms (*μ*g) of each species present in the bird; *w* is the weight of an individual worm; *L* is the helminth body length (*μ*m); *D* is the body diameter (*μ*m), and *c* is the worm count of each species (Andrássy 1956). A summary variable ‘total mass of helminth worms’ was calculated as the sum of *W* values for each species of helminth present in the bird. The data on each species length and diameter used in the calculations in provided in Table S1. In order to investigate the impact of worm colonization on body weight, a linear model was built with the coefficient associated with the variable ‘Presence of helminths’, including bird age, weight, and status (diseased/healthy). The agreement between presence of parasite of each species over subsequent cycles of production was investigated by calculating kappa values. The Bernoulli spatial model was used to identify potential farm clusters of helminth species using SaTScan software (Information Management Services Inc.) using both normal and diseased flocks (Kulldorff 1997). Linear multivariable regression models were built with study variables in order to identify factors associated with burden of each parasite species. The outcomes modelled were: (1) No. of *A. galli* worms (log); (2) No. of *H. gallinarum* worms; and (3) Total No. of cestode worms (log). Variables that were significant at p<0.05 level were left in the final models.

## Results

### Farms and flocks

A total of 120 normal flocks (81 farms, 120 chickens) and 45 ‘diseased’ flocks (45 farms, 90 chickens) were investigated. Diseased flocks were located in a total of five districts. About half (49.7%) of all flocks were located in farms in Thap Muoi (district), whereas 37.0% flocks were in farms in Cao Lanh; the rest were located in Cao Lanh city (5.5%), Lap Vo (4.2%), Tam Nong (2.4%) and Thanh Binh (1.2%).

The descriptive features of chicken farms are shown in Table S2. The median size of normal and diseased flocks was 300 [IQR (interquartile range), 200-505] and 165 birds [IQR 100-300], respectively (Kruskal-Wallis *χ*^2^=27.047; *p*<0.001). The median age of chickens of normal and diseased flocks was 18 [IQR 17-20] and 7 weeks [IQR 4-10], respectively (*χ*^2^=141.97; p<0.001). The median weight of chickens investigated was 1,800g [IQR 1,500-2,200] (normal flocks) and 400.0 g [IQR 185.0-632.5] (diseased flocks) (*χ*^2^=150.78; p <0.001). Chickens in normal flocks were predominantly raised on a soil/earth floor (59.2 %), with a cement floor (23.3%), house on stilts on the ground or water (9.2%) and ‘other’ such as a house with wood, sand, metal, or a metal mesh floor (8.3 %). Anthelminthics had been used in a third (33.3%) of normal flocks over the production cycle. Data on type of floor and anthelminthic use in diseased flocks was not available.

### Prevalence, burden and distribution of gastrointestinal helminths

Over half of all chickens were colonized by gastrointestinal helminths (65/120 normal birds, 54.2% (95 %CI 0.45-0.63); 49/90 diseased birds, 54.4% (95 %CI 0.44-0.65). Nematodes were the most common type of helminth (52.5-54.4% birds colonized), followed by cestodes (15.8-16.7 %), and trematodes (0-1.0%). A total of 8 different species were identified (Table 1). In healthy birds, the greatest prevalence of colonization was: *Heterakis gallinarum* (43.3%) followed by (in decreasing prevalence) *Ascaridia galli* (26.7%) (nematodes), *Raillietina tetragona* (8.3%), *Raillietina cesticillus* (4.2%), and *Hymenolepis* spp. (2.5%) (cestodes). In diseased birds, the highest prevalence of colonization corresponded to *H. gallinarum* (42.2 % birds), *A. galli* (41.1%), and *R. tetragona* (11.1%). The prevalence of colonization with *A. galli* in diseased birds was significantly higher that in healthy birds (*χ*^2^=4.231; p=0.04).

**Table 1:**
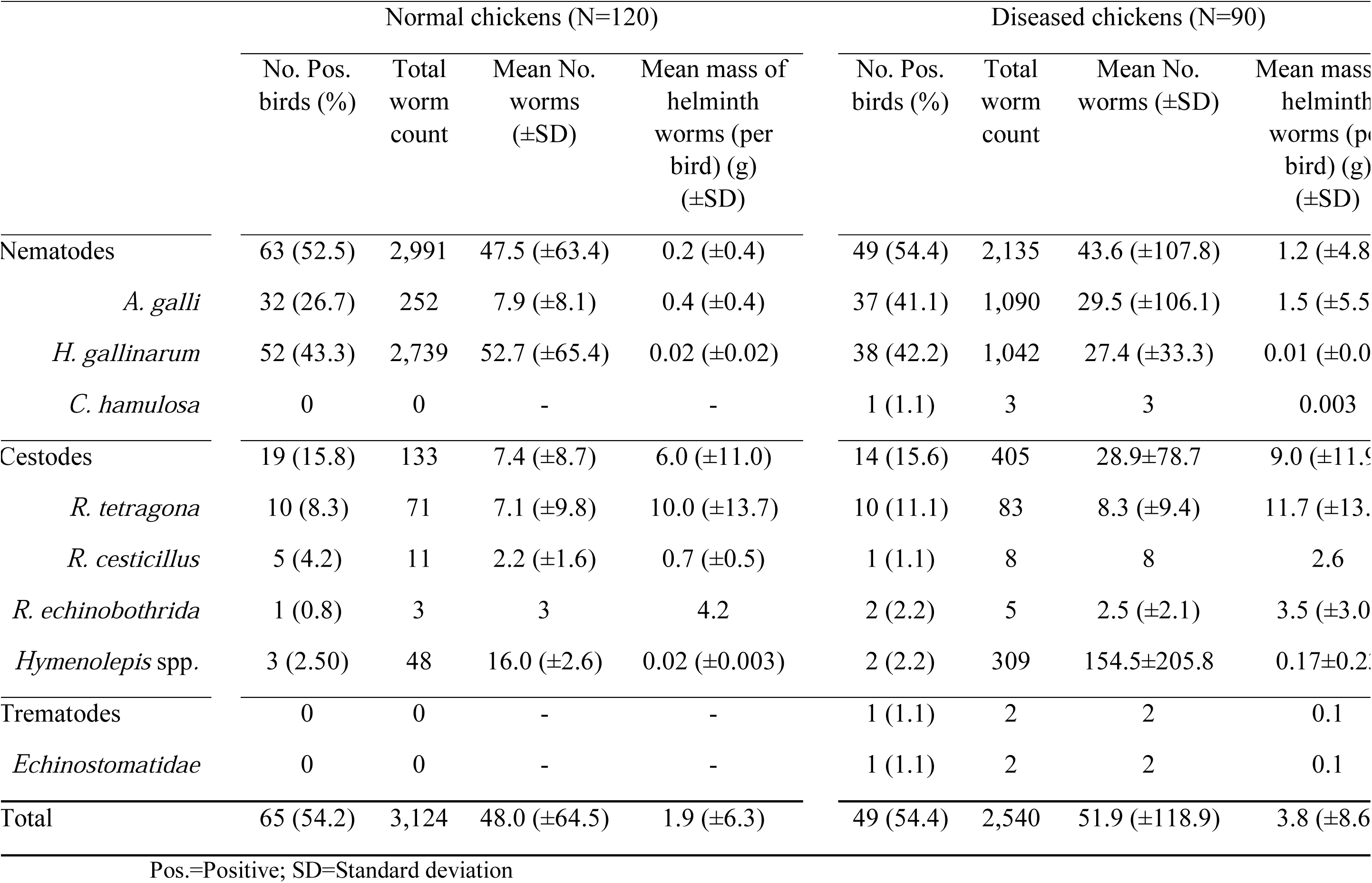
Prevalence (%) of colonization, mean number of helminths of each helminth species, and weight of worms (per bird) among colonized chickens.

Among colonised birds, those that had clinical signs harboured a higher mass of helminth worms than healthy birds (3.8 ±SD 8.6g vs. 1.9 ±6.3g, respectively). This is largely because diseased birds had about a four-fold higher *A. galli* worm count (29.5 ±SD 106.1 worms) compared with healthy birds (7.9 ±SD 8.1 worms). The distribution of different helminth species in colonised birds, stratified by disease status is shown in Figure 2. The counts were most skewed for *H. gallinarum* in normal chickens (median 20, mean 52.7) Conversely, the number of *A. galli* worms, and mass of helminth worms was also considerably more skewed in diseased birds compared with healthy birds. We found a fair to moderate level of agreement between presence of *A. galli* and *H. gallinarum* and cestodes, both in diseased and normal chickens (all p≤0.001). The highest level of agreement was found between colonization with both *A. galli* and *H. gallinarum* in diseased birds (Kappa=0.482; SE 0.104) (Table S3).

**Figure 1:**
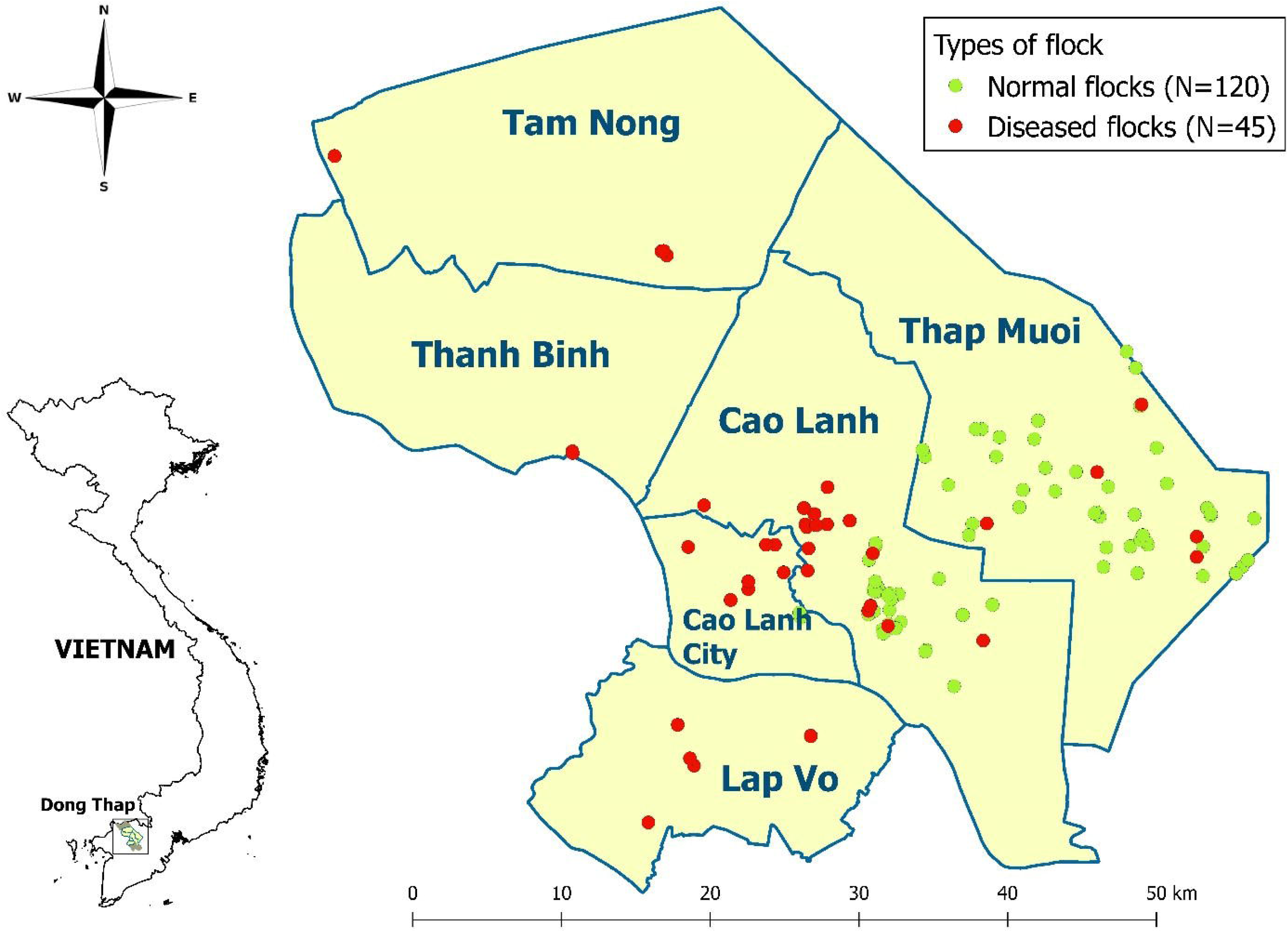
Location of study farms, including farms with ‘normal’ flocks (N=120) and farms with ‘diseased’ flocks (N=45). The names corresponds to districts within Dong Thap province.

**Figure 2:**
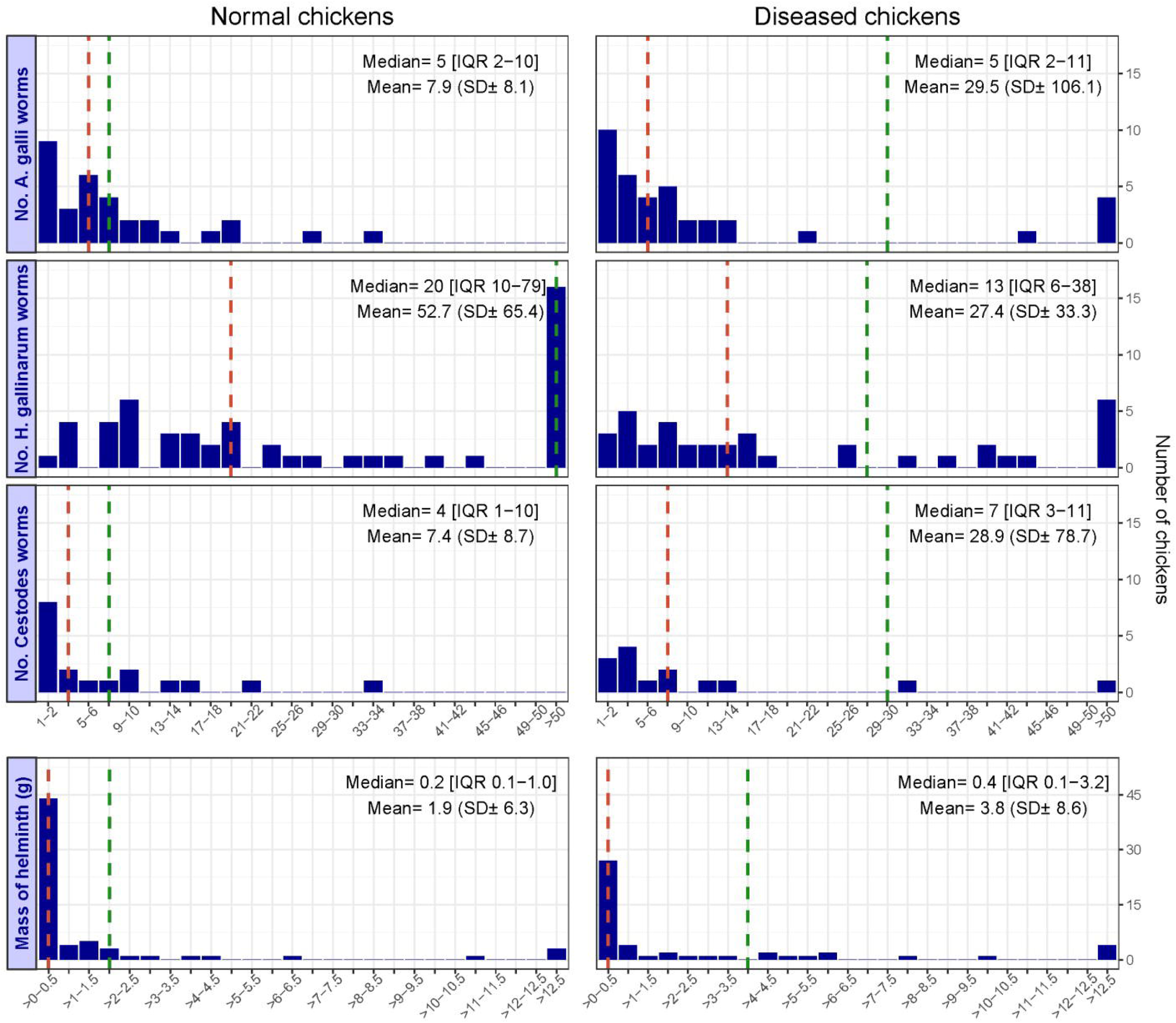
Distribution of chickens by their counts of worms by species in normal and diseased chickens, and mass of helminth worms (in g). The mean (green broken line) and median (red broken line) values are displayed in the graph.

### Prevalence of gastrointestinal helminths and rainfall

The prevalence of chickens colonised with gastrointestinal helminths was 66.7 % (95 %CI 0.57-0.76) and 41.9 % (95 %CI 0.32-0.52%) in the rainy and dry season, respectively (*χ*^2^=12.0; *p*<0.001). Similar differences were seen for all three types of helminth (*A. galli, H. gallinarum* and cestodes) (Figure 3).

**Figure 3:**
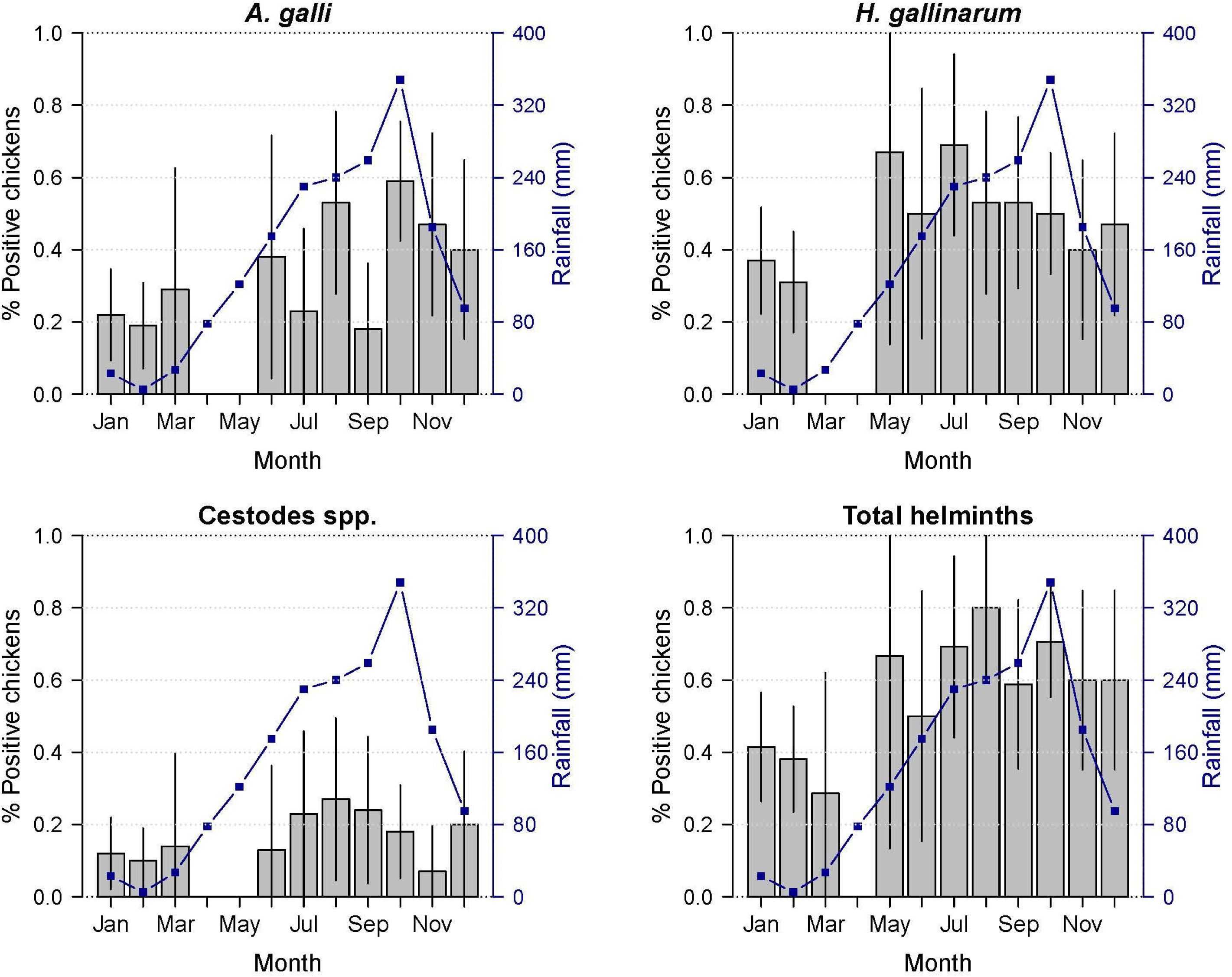
Monthly rainfall (mm) and proportion of helminth-colonized chickens. Vertical lines indicate 95 % confidence intervals around the average prevalence of colonization by month.

### Colonization with helminths over subsequent cycles

In total there were data for 39 flock transitions in 31 normal farms, due to the fact that these farms had more than two cycles. There was no significant correlation between *A. galli* (Kappa=-0.021; Standard Error (SE) 0.151), p=0.445); *H. gallinarum* (Kappa=0.113; SE 0.129; p=0.189) or cestodes (Kappa=- 0.045; SE 0.115; p=0.3488).

### Mass of helminths and chicken body weight

The relationship between helminth infection status and chicken weight and age is displayed in Figure 4. The model predicted that colonized chickens had a body weight lower in 101.5g (95% CI 0-213.2g).

**Figure 4.**
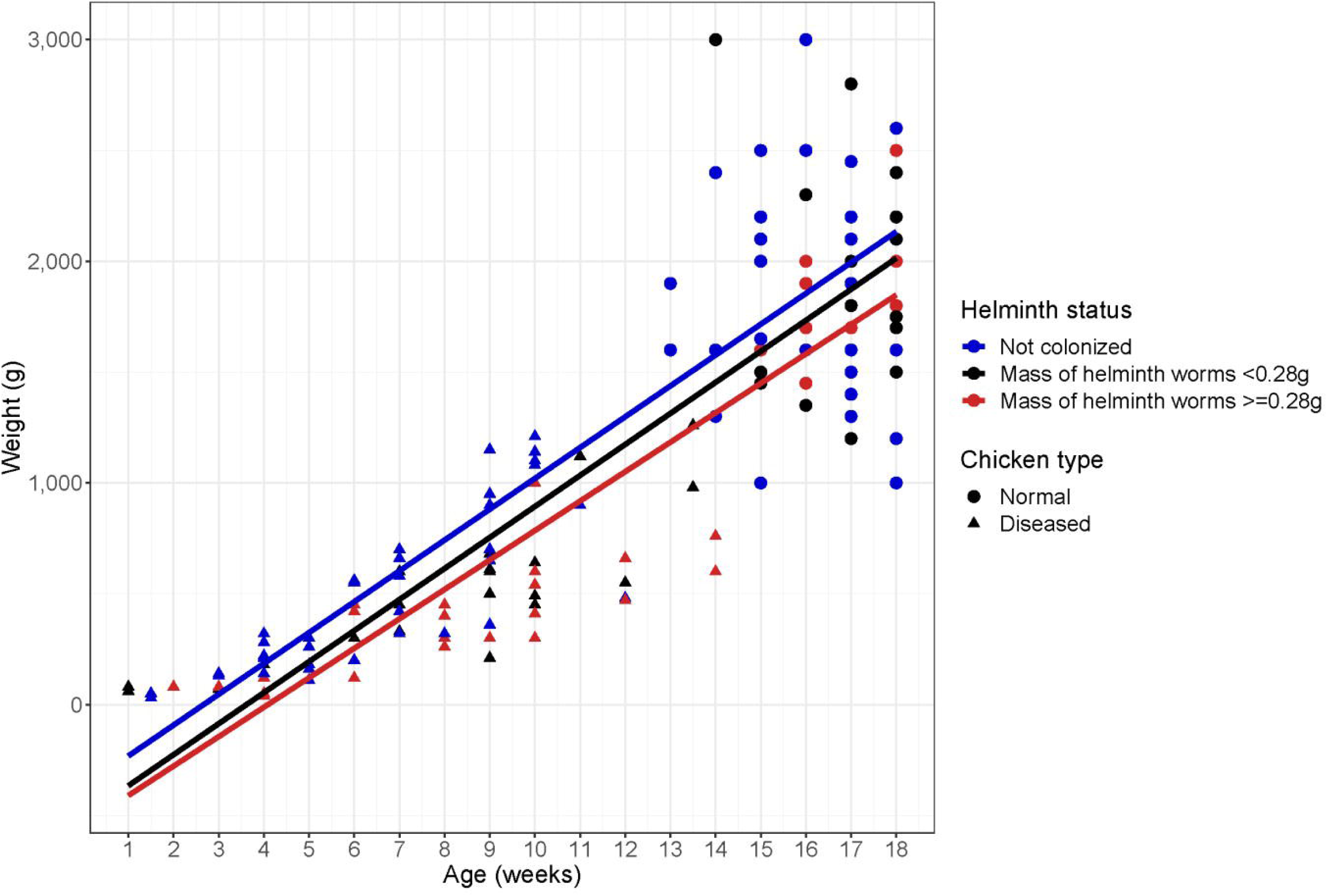
Helminth colonization status, chicken weight and age.

### Use of anthelminthics

A total of 40/120 (33.3%) normal flocks reported using anthelminthics. A total of 9 farmers had administered anthelminthic to their flocks over the five-week period prior to slaughter, and 31 farmers had administered anthelminthic earlier. A total of seven types of anthelminthics had been used: levamisol (42.5% flocks), followed by menbendazol (20.0%), fenbendazol (17.5%), praziquantel (15.0%), ivermectin (7.5%), sufadimethocine (7.5%) and albendazol (2.5%). Figure 5 (supplementary) shows the timing (week) of use of anthelminthics in flocks in relation to the mid-point of the cycle.

**Figure 5.**
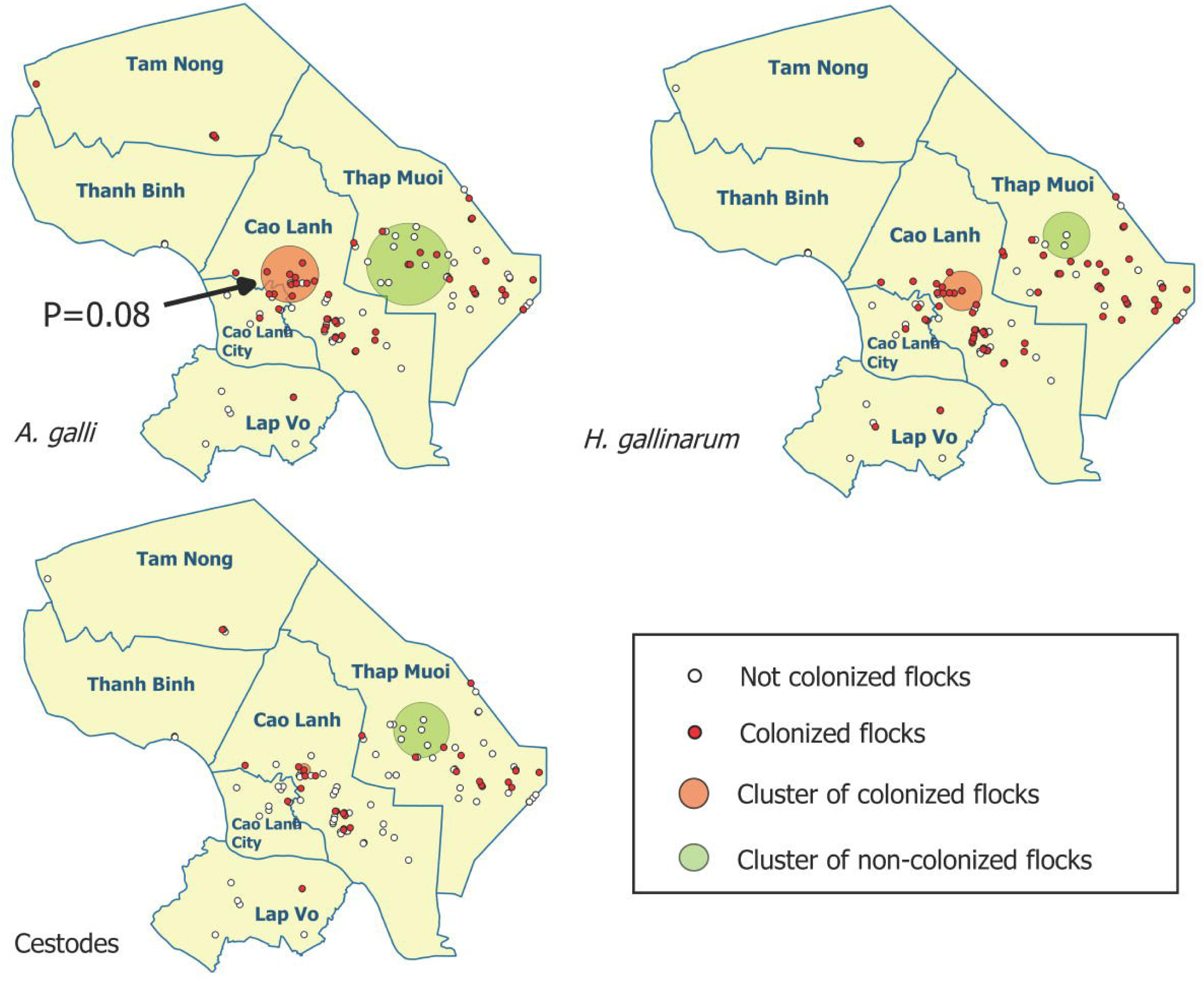
Spatial distribution of flocks colonised and not colonised with *A. galli, H. gallinarum* and cestodes in study flocks. The arrow indicates a significant cluster at p=0.08.

### Risk factor analyses

Only two factors remained significantly associated to the presence of *A. galli* worms: Rainy season (p=0.039) and cement floor (compared with soil, stilts and other types of floor) (borderline significant, p=0.089). Both For *H. gallinarum*, rainy season (p<0.001) and presence of ducks (protective) (p=0.029). For cestodes, cement floor (p=0.009) and presence of ducks (p<0.023) (protective) (Table S4).

### Geographical clustering of helminth

A cluster of low prevalence of colonisation was found for each of the types of parasites (*A. galli, H. gallinarum* and cestodes) in the district of Thap Mui. Clusters of high prevalcne of colonisation was also seen for *A. galli* and *H. gallinarum* in Cao Lanh. However only the cluster in Cao Lanh was significant with p<0.1 (p=0.08) (Figure 5).

## Discussion

We document for the first time the prevalence, burden, and species identity of helminth parasites in healthy and diseased meat chickens raised in commercial, small-scale farms in the Mekong Delta of Vietnam. Our study showed that ∼54% of chickens raised in these systems were colonized by gastrointestinal helminths. Gastrointestinal helminths are not regarded an issue in chicken flocks raised in intensive farming systems with a fast growth cycle (i.e. broilers), and are normally associated with poor biosecurity and deficient terminal cleaning and disinfection (Permin and Hansen 1998). Studies on gastrointestinal helminths performed in other countries in Asia show high (>72%) but variable levels of nematode colonisation in so-called ‘backyard’ and ‘local’ chicken flocks at the end of production, with *H. gallinarum* and *A. galli* being the two most common nematode species (Abdelqader et al. 2008, Alam et al. 2014, Butboonchoo and Wongsawad 2017, Javaregowda et al. 2016, Katoch et al. 2012b, Wuthijaree et al. 2018). The relatively lower prevalence of colonization in birds in our normal flocks compared with published surveys probably reflect the fact that these flocks were all raised in confined conditions most of the time. However, sick chickens were younger and generally came from smaller flocks, but had a similar prevalence of colonization to chickens in normal flocks. Because of this, is is not entirely possible to know to what extent the observed differences reflects an effect on helminths on disease susceptibility, or differences in prevalence associated with the age of the chickens.

In our study *H. gallinarum* was the most prevalent nematode in non-diseased chickens (43% prevalence), and was found at higher prevalence than *A galli* (27%). In diseased chickens, however, levels of colonization with *H. gallinarum* and *A. galli* were comparable in terms of prevalence (41-42%). Given the much higher mass of *A. galli* worms (∼52mg) compared with *H. gallinarum* (∼0.34mg), this resulted in a considerable higher mass of helminth worms, and a clear association between colonization with *A. galli* and disease. It is likely that conditions in farms that favour transmission of respiratory disease are also conducive for helminth colonisation. We tested respiratory samples from sick chickens for *Pasteurella haemolytica* by PCR, and found a greater detection rate of this bacteria among *A. galli*-positive birds than *A. galli*-negative birds (13.5% vs. 1.8%) (personal communication). It has been shown that *A. galli* increases the risk of chickens to outbreaks of fowl cholera (Dahl et al. 2002). An additional negative impact of this parasite is its influence on both humoral and cell-mediated immune responses to Newcastle disease vaccination (Pleidrup, Dalgaard, Norup, Permin, Schou, Skovgaard, Vadekaer, Jungersen, Sorensen and Juul-Madsen 2014). We do not know the vaccination status of these flocks, but they all tested negative to Newcastle disease by PCR. Our normal study flocks (with 100-2,000 chickens) are representative of the ‘emerging’ small commercial meat chicken sector in Vietnam. In this country, only 25% of the ∼230M of chickens produced annually are raised in large scale (‘industrial’) farms (Anon. 2018b). The remaining production includes the backyard farms and small-scale farms. Small-scale commercial farms represent an upgrade from backyard production, since flocks are single-age (i.e. ‘all-in-all-out’) and are mostly raised on commercial feed. These production systems are very much on the increase, a phenomenon going in parallel with an unprecedented increase in demand for poultry meat in the country (Anon. 2018c).

However some flocks may be are at some point allowed to access a rage area (normally adjacent to the chicken house). This practice is likely to contribute to transmission of parasites to one cycle to the other, and may partly explain the observation of ∼16% healthy flocks colonized with cestodes. We did not find however evidence that carry over of infection from one cycle to another was a common issue beyond statistical chance, suggesting that chickens may be colonized when accessing areas outside the chicken pens. The presence of echinstomatids in one flock presenting with disease is related to the ingestion of aquatic snails acting as second intermediate host harbouring the metacercariae. In addition, some farmers are known to supplement their flocks with fresh plants, and snails can be attached to aquatic plants used as suplimentary food.

We empirically demonstrated the economic impact of helminth parasites on chicken bodyweight. A study in India compared the body weight of chickens after 3 months that were treated with anthelminthic and compared with a control (untreated group). The observed difference in bodyweight was ∼385g (Katoch, Yadav, Godara, Khajuria, Borkataki and Sodhi 2012a), which was considerably greater than our study (101.5g) However, chickens in ths study were backyard, and the prevalence of cestodes in that study was much higher.

We found a generally higher prevalence of colonization during the rainy season. Climatic conditions such as rainfall and temperature are known to influence the population dynamics of helminths (Mas-Coma et al. 2008). The Mekong Delta is characterized by high temperature and humidity, a climate highly suitable for the development of parasites outside their final host, facilitating egg embryonation (Tarbiat et al. 2015) (Saunders et al. 2000), as well as the presence of insect intermediate hosts (in the case of cestodes and trematodes) (Abebe et al. 1997).

A surprising finding was the apparently absence of differences in colonization between flocks treated and not treated with anthelmintics. However most farms had used an anthelmintic drug only sporadically, and relatively few farms had applied anthelmnthic in the latter phase of the production cycle. The interpretation of this is unclear, since it can reflect either inadequate preparation of the products (wrong storage, dilution, etc.) or the presence of of anthelmintic resistance against the products used.

Flocks raised on cement floor had unexpectedly a higher prevalence of colonisation with *A. galli* and cestodes compared with those raised on soil/earth floor and stilt houses. We cannot explain these differences, since it is generally assumed that an earth floor contributes to maintain the life cycle of these parasites, since eggs of nematodes can survive for longer periods (Anderson 2000). Stilt houses are likely to limit the risk of infection since faecal material directly falls directly on a minipit underneath the chicken house, or into a pond/canal. The presence of ducks was associated with a lower risk of *H. gallinarum* and cestodes. A possible explanation for the lower prevalence of cestodes in the presence of ducks is their capacity for predating arthropod vectors.

In summary, we characterized levels and burdens of colonization with helminths in chickens raised in emerging, commercial small-scale chicken farming systems in the Mekong Delta of Vietnam. We demonstrated that nematodes account for loss in productivity of such flocks, nematodes being the most abundant type of parasite, and *A. galli* being associated with disease in flocks. We recommend to step up biosecurity and hygiene measures aiming at reducing the prevalence of colonization with helminths. The observed lack of clear evidence of a protective effect from anthelmintic use warrants further investigation.

## Acknowledgements

We would like to thank the staff of the SDAHP-DT for their support with farm sampling and sample processing. This work has been funded by the Wellcome Trust through an Intermediate Clinical Fellowship awarded to Dr. Juan Carrique-Mas (Grant No.110085/Z/15/Z).

## Compliance with ethical standards

This study was a component of the ViParc project, which was granted ethics approval by the Oxford Tropical Research Ethics Committee (OXTREC) (Ref. 5121/16) and by the local authorities (People’s Committed of Dong Thap province) (May 2016). Handling of animals for sampling and euthanasia were conducted in consideration of animal welfare, and followed international guidelines. All participating farmers provided informed consent.

## Conflict of Interest

The authors declare that they have no conflict of interest.

