## Supplementary material for "Characterisation of gastrointestinal helminths and their impact in commercial small-scale chicken flocks in the Mekong Delta of Vietnam": Figure S1

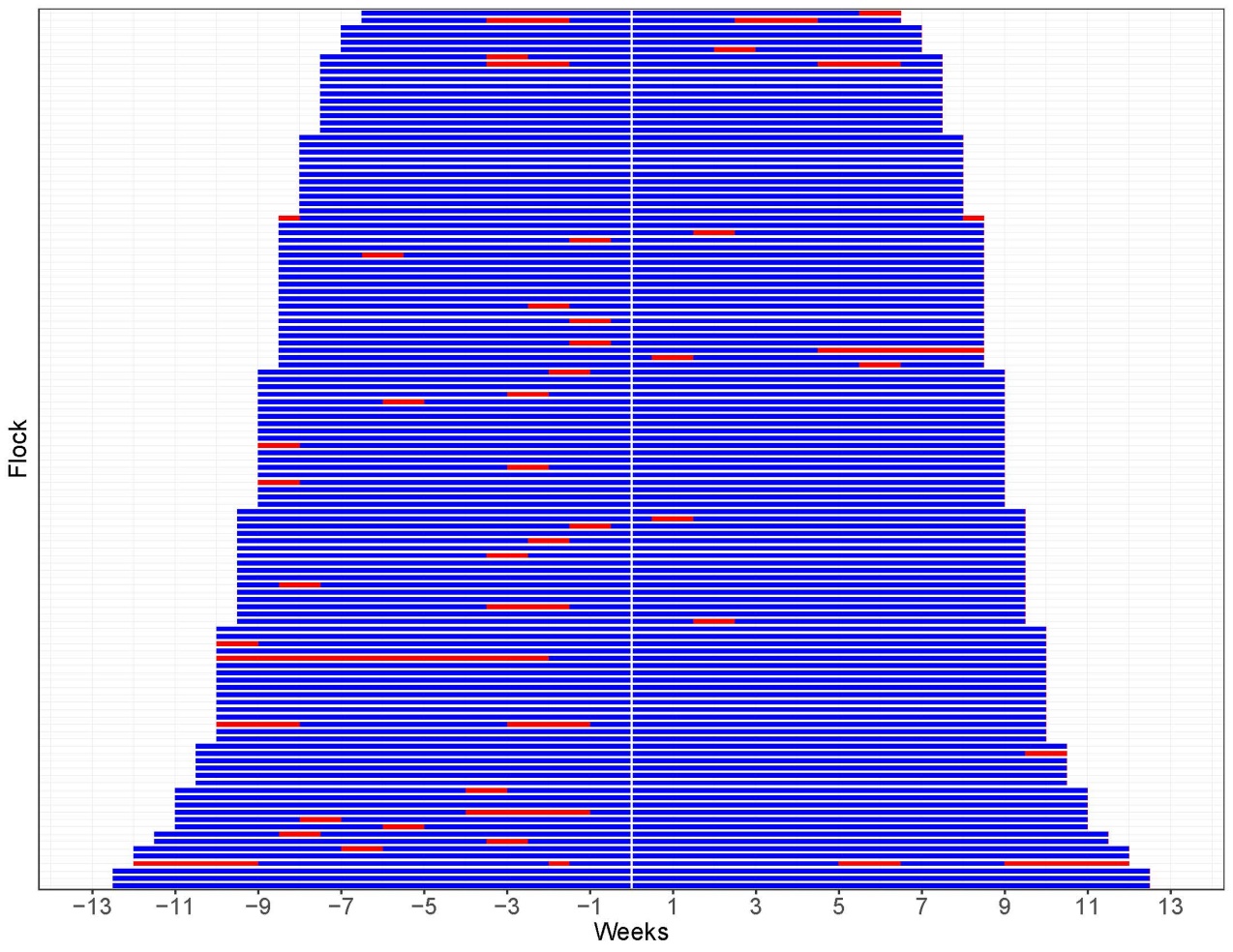
Figure S1: Time of anthelminthic use in 120 normal flocks by week. Flocks are sorted by the duration of the cycle. Blue=Not used anthelminthic. Red=Used anthelminthic.
