## Supplementary material for "Characterisation of gastrointestinal helminths and their impact in commercial small-scale chicken flocks in the Mekong Delta of Vietnam": Table S1

Table S1. Estimated dimensions and mass of worm by species.

| Helminth species | *L* (*µ*m) | *D* (*µ*m) | *w* (mg) | Reference |
| --- | --- | --- | --- | --- |
| *Ascaridia galli* | 69,000 | 1,100 | 52.20 | Morphology and Prevalence of some Helminth Parasites in *Gallus domesticus* from Gurez Valley of Jammu and Kashmir, India |
| *Heterakis gallinarum* | 7,000 | 280 | 0.34 | Morphology and Prevalence of Some Helminth Parasites *in Gallus domesticus* from Gurez Valley of Jammu and Kashmir, India |
| *Cheilospirura hamulosa* | 14,850 | 345 | 1.10 | https://www.ncbi.nlm.nih.gov/pmc/articles/PMC4674620/pdf/JPR2015-569340.pdf |
| *Raillietina tetragona* | 250,000 | 3,000 | 1,406.30 | https://parasitipedia.net/index.php?option=com_content&view=article&id=2588&Itemid=2870 |
| *Raillietina cesticillus* | 130,000 | 2,000 | 325.0 | https://parasitipedia.net/index.php?option=com_content&view=article&id=2588&Itemid=2870 |
| *Raillietina echinobothrida* | 250,000 | 3,000 | 1,406.30 | https://parasitipedia.net/index.php?option=com_content&view=article&id=2588&Itemid=2870 |
| *Hymenolepis* spp*.* | 20,000 | 300 | 1.12 | The Chicken Health Handbook, 2nd Edition: A Complete Guide to Maximizing Flock Health and Dealing with Disease |
| Echinostomatidae | 16,000 | 2,250 | 50.63 | http://www.fao.org/docrep/018/x0583e/x0583e.pdf |

*w*=Estimated weight of an individual helminth worm; *L*=body length (*μ*m); *D*=body diameter (*μ*m).
