## Supplementary material for "Characterisation of gastrointestinal helminths and their impact in commercial small-scale chicken flocks in the Mekong Delta of Vietnam": Table S2

Table S2. Descriptive variables of farms with ‘normal’ (N=120) and ‘diseased’ (N=45) chicken flocks.

| Variable | Level | Normal flocks | | Diseased flocks | |
| --- | --- | --- | --- | --- | --- |
|  |  | **N** | **%** | **N** | **%** |
| Age of farmer (years) | <=45 years | 56 | 46.7% | 24 | 53.3% |
|  | >45 years | 64 | 53.3% | 21 | 46.7% |
| Farmer’s gender | Female | 17 | 14.2% | 6 | 13.3% |
|  | Male | 103 | 85.8% | 39 | 86.7% |
| Highest level of education attained | Primary or secondary | 84 | 70.0% | ND |  |
|  | High school or higher | 36 | 30.0% | ND |  |
| Experience in chicken farming (years) | 0-3 | 67 | 55.8% | 22 | 48.9 % |
|  | >3 | 53 | 44.2% | 23 | 51.1% |
| Type of chicken house floor | Soil | 71 | 59.2% | ND |  |
|  | Cement | 28 | 23.3% | ND |  |
|  | Other | 21 | 17.5% | ND |  |
| Stocking density (chickens/m^2^) | <1 | 28 | 23.3% | 14 | 31.1% |
|  | 1-4 | 59 | 49.2% | 9 | 20.0% |
|  | >4 | 33 | 27.5% | 16 | 35.6% |
|  | ND | 0 | 0.0% | 6 | 13.3% |
| Other chicken flocks in the farm |  | 57 | 47.5% | 9 | 20.0% |
| Ducks in the farm |  | 47 | 39.2% | 13 | 28.9% |
| Muscovy ducks in the farm |  | 28 | 23.3% | 7 | 15.6% |
| Season | Rainy  0 | 69 | 57.5% | 17 | 37.8% |
|  | Dry | 51 | 42.5% | 28 | 62.2% |
| Anthelmintic used* | Never used | 83 | 59.2% | ND |  |
|  | Used >7 weeks before slaughter | 19 | 15.8% | ND |  |
|  | Used 7 weeks before slaughter | 18 | 15.0% | ND |  |

ND=No data
