## Supplementary material for "Characterisation of gastrointestinal helminths and their impact in commercial small-scale chicken flocks in the Mekong Delta of Vietnam": Table S3

Table S3. Comparison between the prevalence of different types of parasite in healthy and diseased birds.

|  | (+)(+) | (+)(-) | (-)(+) | (-)(-) | Kappa value | p-value | Level of agreement |
| --- | --- | --- | --- | --- | --- | --- | --- |
| *Healthy birds* |  |  |  |  |  |  |  |
| AG vs. HG | 21 | 11 | 31 | 57 | 0.254 (0.085) | 0.001 | Fair |
| AG vs. cestodes | 12 | 29 | 6 | 82 | 0.356 (0.086) | 0.001 | Fair |
| HG vs. cestodes | 15 | 37 | 3 | 65 | 0.275 (0.071) | <0.001 | Fair |
| *Diseased birds* |  |  |  |  |  |  |  |
| AG vs. HG | 26 | 11 | 12 | 43 | 0.482 (0.104) | <0.001 | Moderate |
| AG vs. cestodes | 53 | 25 | 2 | 12 | 0.321 (0.085) | <0.001 | Fair |
| HG vs. cestodes | 53 | 25 | 1 | 13 | 0.357 (0.083) | <0.001 | Fair |

AG=*A. galli*; HG=*H. gallinarum*
