## Supplementary material for "Characterisation of gastrointestinal helminths and their impact in commercial small-scale chicken flocks in the Mekong Delta of Vietnam": Table S4

Table S4: Coefficients from linear regression models investigating variables associated with the number of three type of helminth worms (log) in normal chicken flocks. Only variables with p<0.10 in at least one model are shown.

|  | **Model 1**  **(Outcome=*A. galli*)** | | **Model 2**  **(Outcome=*H. gallinarum*)** | | **Model 3**  **(Outcome= Cestodes)** | |
| --- | --- | --- | --- | --- | --- | --- |
|  | Univariable | Multivariable | Univariable | Multivariable | Univariable | Multivariable |
| Farmer’s age (years) (log) | -0.32 (0.29) |  | -0.19 (0.75) |  | 0.36 (0.11) |  |
| Gender (female) | 0.57 (0.02) |  | -0.36 (0.49) |  | 0.30 (0.11) |  |
| Length of cycle (log) | 1.08 (0.07) |  | 1.94 (0.10) |  | 0.47 (0.31) |  |
| House floor  (Ref. Other) |  |  |  |  |  |  |
| *Cement* | 0.41 (0.05) | 0.36 (0.09) | 0.38 (0.37) |  | 0.35 (0.03) | 0.42 (0.01) |
| Density (chickens/m^2^) (log) | 0.05 (0.66) |  | 0.13 (0.56) |  | -0.15 (0.08) |  |
| Ducks in farm | 0.08 (0.66) |  | -0.61 (0.08) | -0.72 (0.029) | -0.25 (0.06) | -0.30 (0.02) |
| Season (Rainy) | 0.38 (0.02) | 0.34 (0.04) | 1.05 (<0.001) | 1.11 (<0.001) | 0.11 (0.39) |  |
| Anthelmintic used  (Ref. Never used) |  |  |  |  |  |  |
| Used >7 weeks before slaughter | 0.07 (0.23) |  | 0.35 (0.46) |  | -0.06 (0.18) |  |
| Used 7 weeks before slaughter | 0.40 (0.24) |  | 0.43 (0.47) |  | -0.12 (0.18) |  |
